# Establishment of an *in vitro* culture system to study the developmental biology (growth, mating and nodule formation) of *Onchocerca volvulus* with implications for anti-*onchocerca* drug discovery and screening

**DOI:** 10.1101/2020.06.25.170746

**Authors:** Narcisse Victor T. Gandjui, Abdel Jelil Njouendou, Eric Njih Gemeg, Fanny Fri Fombad, Manuel Ritter, Chi Anizette Kien, Valerine C. Chunda, Jerome Fru, Mathias E. Esum, Marc P. Hübner, Peter A. Enyong, Achim Hoerauf, Samuel Wanji

## Abstract

**Background:** Infections with *Onchocerca volvulus* nematodes remain a threat in Sub-Saharan Africa after two decades of ivermectin mass drug administration. Despite this effort, there is still an urgent need for understanding the parasite biology, especially mating behaviour and nodule formation, as well as development of more potent drugs that can clear the developmental (L3, L4, L5) and adult stages of the parasite and inhibit parasite’s reproductive and behavioural pattern.

**Methodology/Principal Findings:** Prior to culture, freshly harvested *O. volvulus* L3 larvae from dissected *Simulium* were purified by centrifugation using a 30% Percoll solution to eliminate fly tissue debris and contaminants. Parasites were cultured in both cell-free and cell-based co-culture systems, and monitored daily by microscopic visual inspection. Exhausted culture medium was replenished every 2–3 days. The cell-free culture system supported the viability and motility of *O. volvulus* larvae for up to 84 days (DMEM–10%NCS), while the co-culture system (DMEM–10%FBS–LLC-MK_2_) extended the worm survival period to 315 days. Co-culture systems alone promoted the two consecutive parasite moults (L3 to L4 and L4 to L5) with highest moulting rates observed in DMEM–10%FBS–LLC-MK_2_ (69.2±30 %), while no moult was observed in DMEM–10%NCS–LEC condition. *O. volvulus* adult worms mating and even mating competitions were observed in DMEM–10% FBS –LLC-MK_2_ co-culture system. Early nodulogenesis was observed in both DMEM–10% FBS–LLC-MK_2_ and DMEM– 10%NCS–LLC-MK_2_ systems.

**Conclusions/Significance:** The present study describes an *in vitro* system in which *O. volvulus* L3 larvae can be maintained in culture leading to the development of reproductive adult stages. Thus, this platform gives potential for the investigation of mating, mating competition and early stage of nodulogenesis of *O. volvulus* adult worms that can be used as additional targets for onchocercacidal drug screening.

**Author summary:** River blindness affects people living in mostly remote and underserved rural communities in some of the poorest areas of the world. Although significant efforts have been achieved towards the reduction of disease morbidity, onchocerciasis still affect million of people in Sub-Saharan Africa. The current control strategy is the annual mass administration of ivermectin which have accumulated several drawbacks overtime: as the sole microfilaricidal action of the drug, very long treatment period (15-17 years) and reports of ivermectin losing its efficacy; Therefore, raising the urgent need for new onchocercacidal molecules. Our study has established an *in vitro* platform capable of supporting the growth and development of all developmental stages of *O. volvulus* (L3 infective stage, L4, L5 and adult worms), moreover the platform provided more insight on *O. volvulus* adult worms reproductive and behavioural pattern. Our findings provide more avenues for mass production of different parasite stages, the investigation of parasite developmental biology and the identification of targets for drug discovery against different phases of development of this filaria parasite

## Introduction

Onchocerciasis is the second major cause of infectious blindness and a major public health problem in many parts of the world [1]. The disease is known as river blindness, because its causative agent *Onchocerca volvulus* is transmitted by *Simulium* (blackfly) vectors, which breed in fast-flowing rivers. Onchocerciasis is endemic in 37 countries in West, East and Central Africa, the Arabian Peninsula and parts of South and Central America. Globally, about 90 million people are at risk of contracting the disease in endemic areas, with 99% of cases occurring in Sub-Saharan Africa, of which more than 17 million are estimated to be infected and 270,000 are permanently blind as complication [2]. In the past decades, two programmes were implemented to control onchocerciasis. Initially, the Onchocerciasis Control Programme (OCP) focused on elimination of the *Simulium* vector using DDT and larvicides. This vector control programme was later supplemented with ivermectin distribution that has helped many millions of persons to live free of disease [3]. Mass drug administration (MDA) of ivermectin through the African programme for Onchocerciasis control (APOC) has been the principal strategy for onchocerciasis control after OCP in West Africa.

Despite more than two decades of MDA campaign with ivermectin, the disease still persists mainly in sub-Saharan Africa because of several reasons. Ivermectin is solely microfilaricidal with temporal embryo-static effects on adult worms and thus has to be given once or twice per year for the life-span of the adult worms, which is 16–18 years. In addition, the sub-optimal response to ivermectin in some regions has led to persistent transmission [4–6]. Therefore, there is a need for the development of improved drugs that might not only kill *O. volvulus* microfilariae but also other life-cycle stages (L3, L4, L5 and adult worms). Such new candidates are required to reach the United Nations Sustainable Development Goals to eliminate onchocerciasis by 2030 in the majority of endemic countries. However, the advancement of research towards the development of new therapeutics is hindered by the availability of a suitable *in vitro* culture system, where parasite stages such as infective larvae (L3) can be maintained and developed into adults. An artificial system which can mimic the human host micro-environment and support the growth and development of *O. volvulus* parasites from the infective stage larvae (L3) to adults would be ideal.

Literally, little is known about the time course of *O. volvulus* parasite developmental process, mating behaviour and nodule formation in the human host. Nevertheless, it is reported that moulting from L3 to L4 occurs within a week (3–7 days) [7], while the L4 to L5 moult is estimated to occur after 2 months [8]. Early L5 are considered young adults, and at this stage the worms have partially developed gonads [9]. In the cattle model, it takes 279–532 days post infection for the closely related *O. ochengi* parasite to develop into fully mature and fertile adult worms capable of releasing microfilariae (mf), the worms’ offspring [10]. More than 400 days post infection is required for the same achievement in a chimpanzee model for *O. volvulus* [8, 11].

Many non-conclusive attempts have been carried out to develop the complete life cycle of *O. volvulus* in artificial *in vitro* systems, though these contributions have been considered as important milestones towards achievement this ultimate goal [7, 9, 12, 13, 14, 15, 18, 21, 22, 28, 32, 35, 43]. Serum/cell free systems have been used in several studies to culture filarial parasites and reports have highlighted maintenance of full viable parasites for a week [12–22]. Improvement of the culture conditions have been achieved by supplementing the basic culture media used with serum or other culture ingredients (fatty acids and complex lipids formulation). The serum-based culture systems have been reported to support parasite longevity *in vitro* and cuticle casting [21, 23–32]. Due to the inconsistency of serum composition, serum-free culture systems and co-culture systems using eukaryotic cells as feeder layers have been found successful [33–43]. Moreover, feeder cells were already shown to be crucial for *in vitro* cultivation and growth of *O. volvulus* [44]. From our previous observations on the *in vitro* growth and development of the filarial nematode *Loa loa* [42], among the three most used supplements (feeder layer, serum, and basic culture medium) for filarial parasites *in vitro* culture, feeder cells were classified as top most important requirement followed by the serum type and finally the nature of the basic culture medium. Summarily, the advancement of research towards the development of a suitable *in vitro* culture system for filarial parasites has highlighted the complexity of their requirement in terms of nutritional needs for growth and moulting from one stage to another. This study aimed at identifying suitable *in vitro* culture requirements, which could support the maintenance and promote the growth and development of the human parasite *Onchocerca volvulus* from its infective L3 larvae stage to reproductive adult worms. Such an *in vitro* culture system will contribute to the experimental production of subsequent parasite stages (L4, L5 and adults of *Onchocerca volvulus*), enable investigations on the reproductive behaviour as well as nodule formation that could be used for further understanding of the parasite biology and identification of novel therapeutic drug targets against onchocerciasis.

## Methods

### Ethical statement

Ethical clearance was obtained from the National Institutional Review board, Yaoundé (N^0^ 2018/06/1057/CE/CNERSH/SP) after approval of the protocol. Prior to recruitment, the nature and objectives of the study were explained to potential participants and those who agreed to take part in the study signed a consent form. Special consideration was taken to minimize any health risks of the participant. They were followed-up for ivermectin treatment at the end of the study during the normal MDA period. Their participation was strictly voluntary and their documents were given a code for confidentiality.

### Determination of *O. volvulus* microfilarial load in skin biopsies of volunteers prior to *Simulium* engorgement

Participants examined were from the Meme drainage basin (overall Community Microfilarial Loads (CMFL) = 5.2 microfilariae / skin snip) and microfilarial load was determined as described by Wanji *et al.* [45]. Briefly, after the clinical examination, two skin biopsies from the posterior iliac crest were taken using a 2 mm corneo-scleral punch (CT 016 Everhards 2218–15 C, Germany). The skin samples from each participant were placed in two separate wells of a microtitre plate containing 2 drops of sterile normal saline. The corresponding well numbers were reflected on the participant’s form. The plates were sealed with parafilm to prevent any spill over or evaporation and incubated at room temperature for 24 hours. All emerged microfilariae were counted using an inverted microscope (Motic AE21) at 10x magnification and expressed per skin snip. Two participants were enrolled in the study that had average microfilariae load of 50 and 65 microfilariae/skin snip respectively.

### Collection of engorged *Simulium* flies

Flies were collected along the banks of a fast-flowing river at Mile 16 Bolifamba (South West region – Cameroon). The fly collection team was composed of two trained individuals, one working from 07:00 AM to 12:00 Noon and the other from 12:00 Noon to 18:00 PM for 5 consecutive days. Female blood-seeking *Simulium* flies were allowed to land on exposed legs of the microfilaridemic donor, where they were allowed to blood-feed and then captured using *Simulium* rearing tubes. Captured-engorged *Simulium* were then transported to the laboratory insectarium and maintained for 10 days for the development of *O. volvulus* infective L3 larvae.

### Laboratory maintenance of engorged Simulium

Blood-fed *Simulium* were maintained in captivity under controlled experimental conditions as described by [58] for 10 days to allow ingested microfilariae to mature into infective stage larvae (L3). Briefly, captive flies were fed on 15% sucrose solution soaked in cotton wool and maintained at 23–28°C and 79–80% relative humidity.

### Dissection of flies, isolation and purification of *O. volvulus* L3

After 10 days of rearing, the flies were dissected in Petri dishes (CytoOne, UK) containing RPMI 1640 medium (Sigma-Aldrich, St Louis, USA). The head, thorax and abdomen were separated and teased apart in three different dishes. Fly tissues were incubated for 20 min to allow L3 larvae to migrate out of the tissue. A sterile pipette was used to pick the larvae and pooled in a shallow convex glass dish [46]. The worms were transferred into 15 ml centrifuge tubes (Corning, Kennebunk-ME, USA) for purification. Only L3 harvested from the head (where more mature larvae are expected to be found) were used in this study. The L3 were washed using a Percoll® (GE Healthcare, Pharmacia, Uppsala, Sweden) centrifugation technique as described by Zofou *et al.* [42]. In summary, the L3 suspension concentrated in less than 1 ml RPMI was slowly layered on the surface of a 15 ml tube containing stock iso-osmotic Percoll® and centrifuged (Humax 14k human, Germany) at 68 *x g* for 10 min. The process was repeated to remove microbial contaminants. At the end, the L3 were washed twice with RPMI-1640 by centrifugation at 239 *x g* for 10 min to remove Percoll® remnant.

### Preparation of feeder cells and pre-conditioning in culture plates

Monkey kidney cells (LLC-MK_2_), mouse lung embryonic cells (LEC), human embryonic kidney cells (HEK-293) and human hepatic cells (HC-04) were provided by the American Type Culture Collection (ATCC, Manassas, Virginia, USA). Each of these feeder cells were cultured in flasks at 37 °C in a humidified CO_2_ incubator (Sheldon Mfg. Inch, Cornelius, OR, USA) at 5% CO_2_ until the cell layer became fully confluent. For new inoculations and other cell manipulations, cells were dislodged with trypsin solution (25%) containing EDTA and kept at 37 °C for less than 30 minutes. The cell suspension was centrifuged at 239 *x g* for 10 min, the supernatant discarded, and the pellet re-suspended and diluted to 10^5^ cells/ml. Aliquots (100 μl) of cell suspensions were plated into each well of a 48-well flat bottom culture plate and kept in the incubator for cells to become fully confluent prior to be used for parasite maintenance in co-culture systems.

### *In vitro* culture of *O. volvulus* larvae

Harvested L3 from different batches of dissected flies were mixed and pooled to obtain parasites culture material. Two sera supplements were used separately at 10 % concentrations each: fetal bovine serum (Sigma-Aldrich, St Louis, USA) and newborn calf serum (Sigma-Aldrich, Berlin, Germany). Five basic media were used: RPMI-1640, IMDM, NCTC-135, MEM (Sigma-Aldrich, St Louis, USA), and DMEM (Gibco Life Technologies, Cergy-Pontoise, France). Penicillin-Streptomycin-Neomycin (PSN, 2 %) was used as antibiotic and Amphotericin B (2.5 μg/ml) as antifungal. Flat bottom culture plates (48-well) with lids (Corning, Kennebunk, ME, USA) were loaded as follows: For the co-culture systems, parasites (range 8–13 L3) in 1200 μL of the studied medium (basic culture medium + 10 % serum) were loaded into a feeder-cell type pre-conditioned plate while in cell-free systems, they were loaded into empty wells. Five batches of infective L3 larvae were used throughout this study and each experimental culture system was carried out in quadruplet wells.

### Assessment of parasite viability

The viability of the parasites was assessed daily, by visual inspection (by two individuals) under an inverted microscope until movement ceded. Their motility was scored on a 4-point scale as described in [47]. Briefly, score 0, no movement or immotile, score 1, intermittent shaking of head and tail, score 2, sluggish (shaking of the whole worm on a spot), score 3, vigorous movement (shaking of the whole worm and migration from one spot to another) was considered.

### Parasite long term *in vitro* maintenance strategy

To achieve long term maintenance, 800 μL exhausted culture medium was removed of each well and replaced with the same volume of fresh culture medium every 2–3 days. Additionally, cultured parasites were transferred from one culture plate (old) to another (new) either when feeder cell growth overshadowed parasites motility scoring (parasite entanglement within overgrown cell) with the following cell lines (HC-04, LEC and HEK-293) every 2 weeks or when feeder cells (LLC-MK_2_) underwent apoptosis every 7 weeks.

### Data processing and analysis

Three variables were used to assess the viability, growth and development of the parasites (mean motility, moulting rate and parasite stage morphometry). Raw data were saved in a spreadsheet and using the above described 4-point scale, the percentage (%) of motility was calculated according to the following formula:

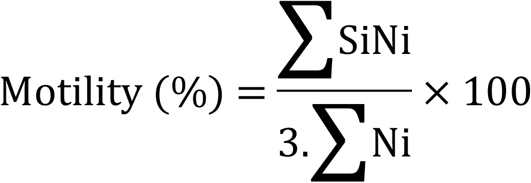

where Si is the score of point scale i and Ni is the total number of worms at a point scale i [47].

Filarial parasite moulting is one of the key phenomena providing clear evidence of worm growth and development. Moulted worms display casted cuticles and morphological changes. The moulting rate was calculated as previously described [41, 42]:

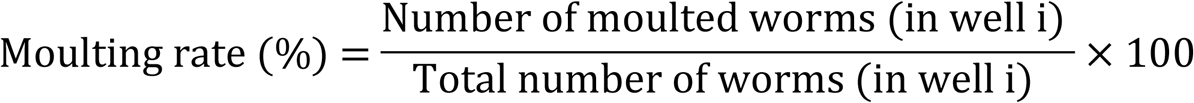

With respect to parasite morphometry, photographs of *O. volvulus* at different stages were recorded under an inverted microscope (OPTIKA, Ponteranica, Italy) and their length was determined using the OPTIKA IS view software Version. 2.0 (Ponteranica, Italy). ImageJ 1.52 software (NIH, USA) was used to generate scale bars of displayed photographs.

GraphPad Prism 8 software (GraphPad, San Diego, USA) was used to generate mean motility, moulting rate and morphometry graphs. Results of replicates were expressed as mean ± standard deviation (SD) for the following variable (motility and moulting) while median was used to summarise morphometric parameters of the parasite. The Kruskal-Wallis one-way analysis test was used to assess differences in motility, moulting rate and worm’s morphometry between sets of studied culture systems. Dunn's *post-hoc* test was applied for pairwise multiple comparisons of the ranked data.

Factors that promoted parasite survival were identified using the multiple linear regression. The general linear model (GLM) was built using the hierarchical stepwise method. A total of 4 blocks were achieved with the 5 factors (incubation time, presence or absence of feeder cells, basic medium, serum) and those that contributed significantly to the improvement of the model were identified based on the F-statistics and the adjusted R-square. The incubation time was treated as a metric factor. Dichotomous variables such as the presence of monkey kidney cells were coded using binary figures. For each nominal factor (Basic culture media, serum), sets of dummy variables were created and compared to one of the categories defined as reference. While RPMI-1640 was used as a reference against DMEM, IMDM, MEM and NCTC. FBS serum was used as a reference against NCS.

The passage of *O. volvulus* larvae from the third (L3) to the fourth (L4) stages then to the fifth (L5) stage was further considered the second target product profile in assessing the suitability of the culture systems tested. Finally, the model was used to predict T_20_ and T_10_ values (Days), defined as the duration (incubation time) at which 20 and 10 % of the worms were still active respectively (score 3). For all statistical comparisons, the *p*-values below 5% were evidence for rejecting null hypothesizes.

## Results

Purified *O. volvulus* infective larvae were cultured in two distinct systems: the cell-free culture system and the cell-based co-culture system. The first step consisted at evaluating the potential of the cell – free systems (combination of each of the five basic culture media and a single concentration of either of the two sera) on the viability, growth and development of *O. volvulus* larvae. Secondly, to subject the best cell – free culture setting on four different mammalian cell lines in order to evaluate the beneficial effect of co – culture with feeder layers (co – culture systems).

### Evaluation of cell-free culture systems on the growth and development of *O. volvulus* larvae

The cell-free system was made of the combination of each of the five basic culture media supplemented with 10 % of any of the two sera (NCS and FBS).

With respect to the cell-free system, the various study culture settings sustained *O. volvulus* larvae viability for a maximum of 84 days. Complete inactivity of all larvae was recorded in culture combinations IMDM – 10 % FBS and NCTC135 – 10 % FBS after 54 days and in DMEM – 10 % NCS and IMDM – 10 % NCS after 84 days. Generally, freshly dissected and cultured *O. volvulus* infective L3 larvae were not vigorously active (motility score = 2, sluggish). Their motility significantly increased from day 3 to day 5 (motility score = 3, vigorously active) when the L3 stage larvae casted their cuticles to become L4. The parasite motility waned in all culture conditions tested after around 45 days of culture and according to the *in vitro* culture combination (Medium-Serum) all parasites were immobile at day 54 to 84 (Fig 1).

**Fig 1.**
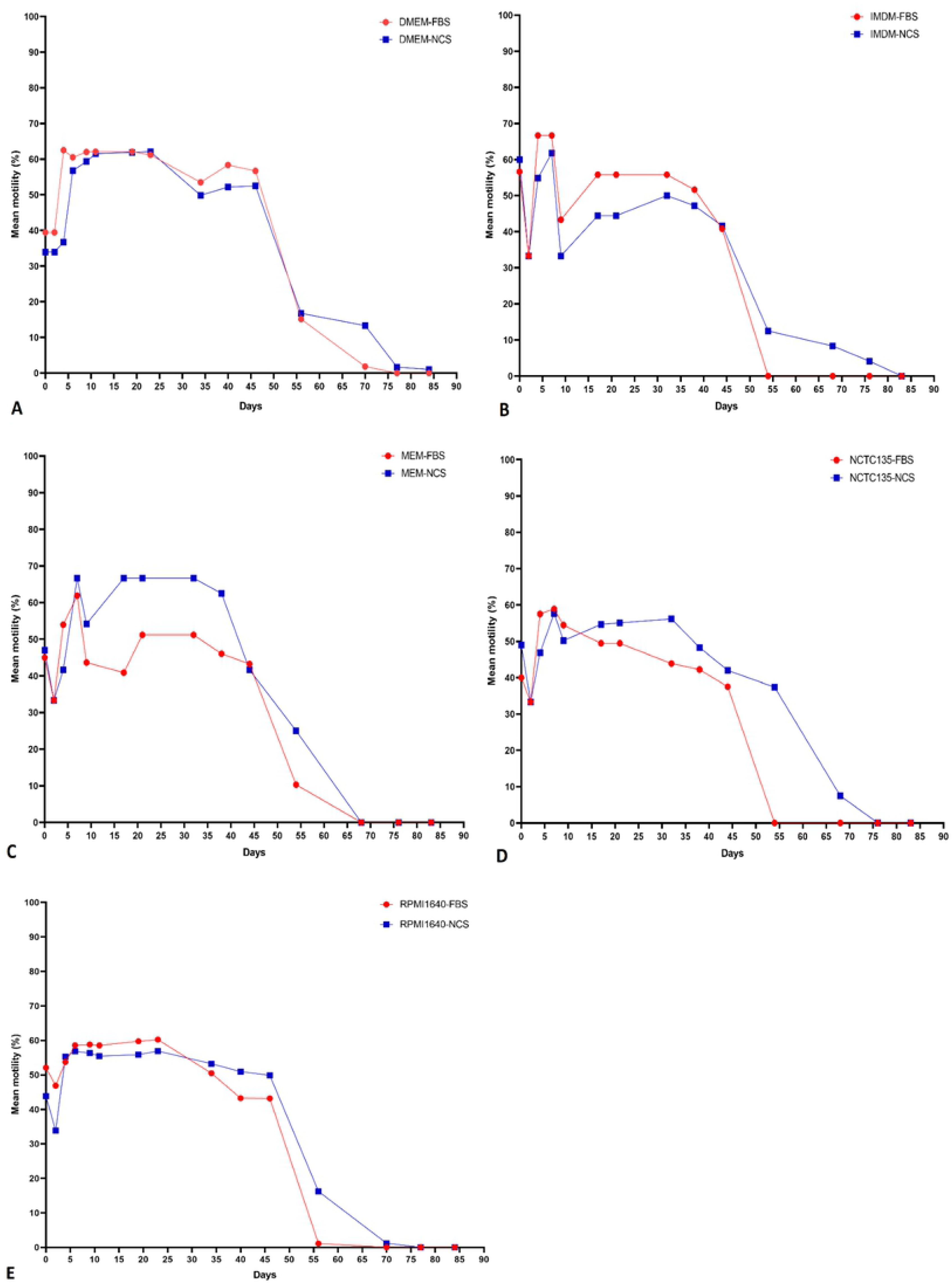
Motility pattern of *O. volvulus* (from L3 to L4) in the cell-free culture systems. **A** DMEM - 10 % FBS/NCS. **B** IMDM - 10 % FBS/NCS. **C** MEM - 10 % FBS/NCS. **D** NCTC 135 - 10 % FBS/NCS. **E** RPMI1640 - 10 % FBS/NCS. (Results were pooled from different independent experiments, n =3 and each experimental setting conducted in quadruplets).

*Onchocerca volvulus* moult was also used as an indicator to assess larvae development *in vitro.* The moult profile of *O. volvulus* larvae *in vitro* was culture system dependent and with respect to cell-free systems, only the first parasite molt (L3 to L4 = M1) was observed. The cuticle of L3 larvae was casted within 3–5 days of culture and the moulting rate ranged from 0 % (MEM – 10 % NCS) to 92 % (DMEM – 10 % NCS). The culture system that best supported *O. volvulus* L3 moult was DMEM – 10% NCS, although no statistically significant difference was observed as compared to DMEM – 10 % FBS, RPMI – 10 % NCS, RPMI – 10 % FBS, NCTC 135 – 10 % FBS and IMDM – 10 % FBS (Fig 2).

**Fig 2.**
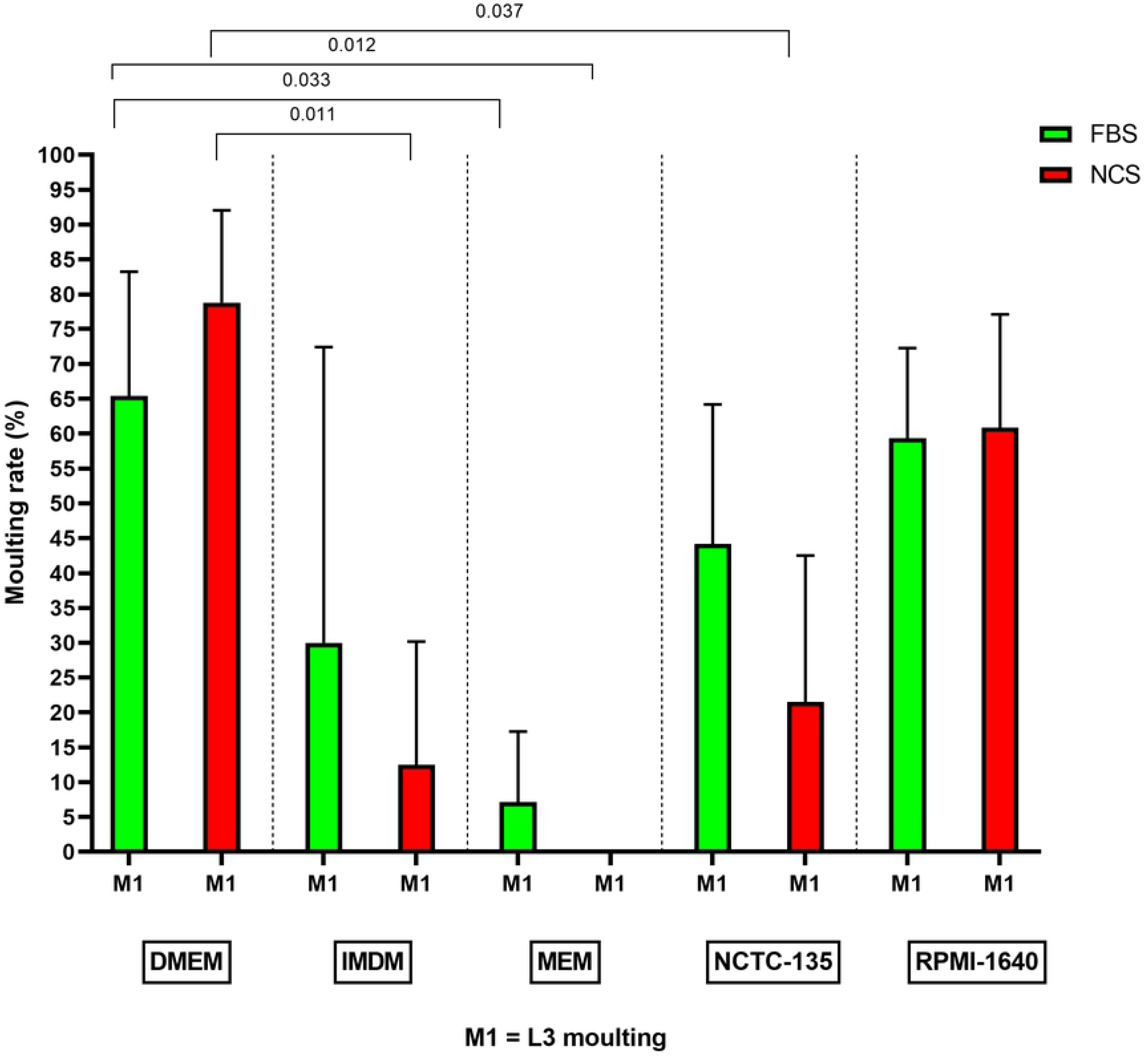
Influence of cell-free culture systems on *O. volvulus* larvae moults *in vitro.* (Results were pooled from different independent experiments [n =3] and each experimental setting conducted in quadruplets).

In cell – free systems, 3 – 5 days were required for *O. volvulus* infective larvae to moult from L3 to L4. Preliminary values of L3 and L4 length and width showed highly overlapping data with no significance, therefore focus was drifted towards comparing L4 and L5 values. In addition, no proof of parasite maturity was recorded since all parasites that succeeded to achieve their first moults (from L3 to L4) failed to undergo the second moult (L4 to L5). The absence of mammalian cells in this system hindered complete parasite growth and development. In summary, the cell – free system supported parasite moult into L4 larvae and viability for up to 84 days, but was below the timeframe necessary to allow full parasite development. Thus, also no cellular mass forming around *O. volvulus* worms was observed in cell – free systems.

### Evaluation of cell-based co-cultured systems on the growth and development of *O. volvulus* larvae

Since larvae did not become adults and stopped their motility by day 84 at the latest in the cell-free culture systems at stage 4, we next evaluated the combination of the best cell – free system (DMEM - NCS/FBS) with each of the four mammalian cell lines (LLC-MK_2_, HC-04, HEK-293 and LEC) in order to improve parasite motility and viability as well as moulting.

Interestingly, *O. volvulus* larvae survived for a longer period of time (up to 315 days) in cell-based co-culture systems (Fig 3). Thus, the culture of *O. volvulus* larvae on feeder cells increased their longevity by 3.75 folds as compare to those cultured in cell-free systems. As similarly observed in cell-free systems, cultured *O. volvulus* infective L3 larvae from freshly dissected *Simulium* flies did not displayed vigorous activity (motility score = 2, sluggish). The infective larvae (L3) motility significantly increased as from day 3 to 7 (motility score = 3, vigorously active) when L3 stages casted out their cuticles to become L4 larvae. The L4 larvae motility remained rectilinear till day 48 when the first L4 to L5 moult was observed. As from day 48, except for larvae cultured in DMEM – 10 % NCS - HC-04, L5 larvae motility started dropping and movement completely ceded in larvae cultured in DMEM LEC 10% NCS after 103 days and in larvae cultured in DMEM HEK 10% NCS after 162 days. The sole L5 larvae motility amplification was observed in DMEM – 10 % NCS - HC-04 culture system with more than 90 % of parasite being very active until 125 days of culture. The motility of this group started dropping afterwards. At day 233 larvae cultured in DMEM – 10 % NCS - HC-04, DMEM – 10 % NCS LLCMK2 and DMEM – 10 % FBS LLCMK2 still showed a motility of 10-20% (Fig 3).

**Fig 3.**
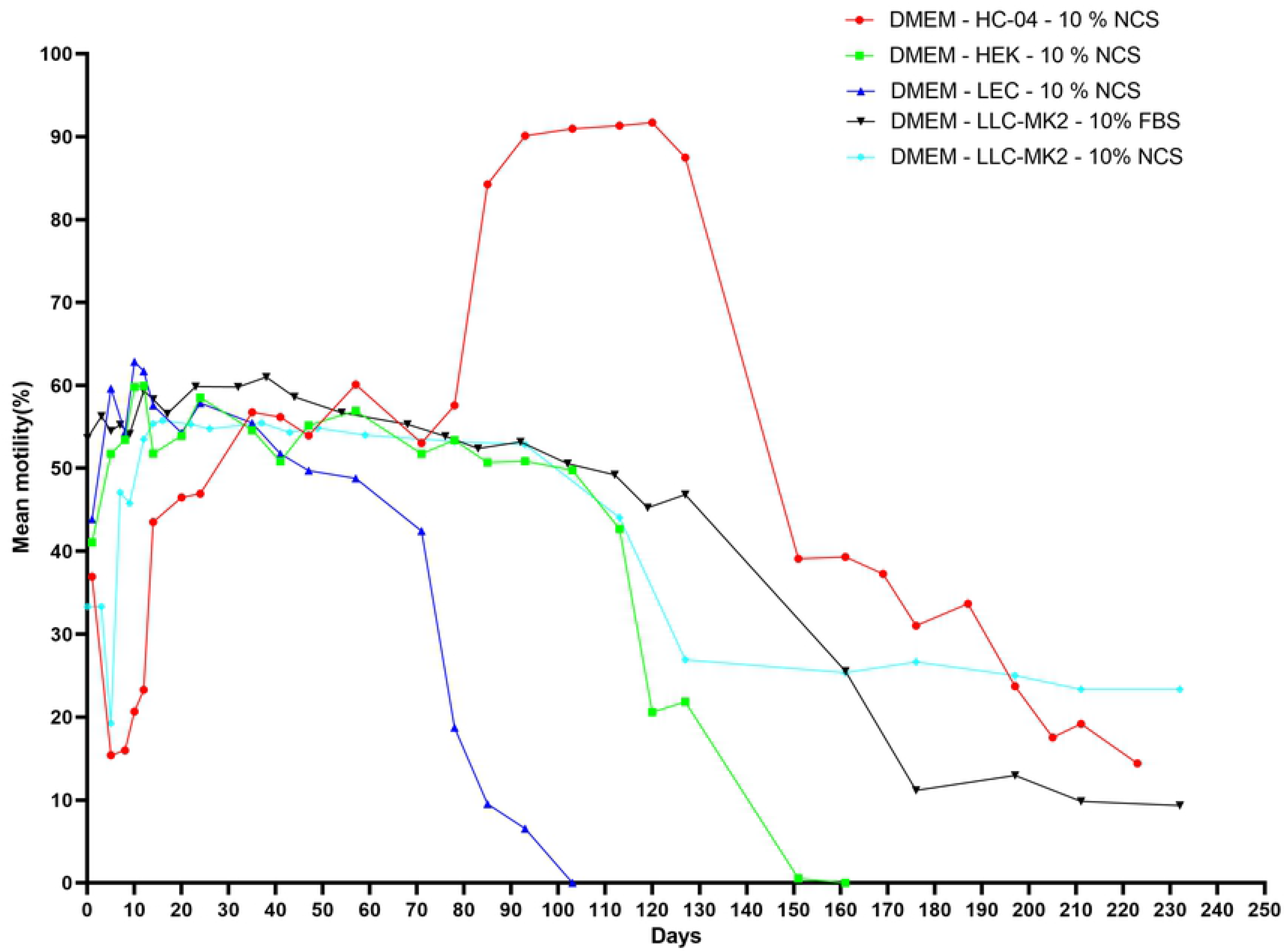
Motility pattern of *O. volvulus* (from L3 - L4 - L5) in the DMEM cell-based co-culture systems supplemented with 10 % FBS and 10 % NCS. (Results were pooled from different independent experiments, n =3 and each experimental setting conducted in quadruplets).

In contrast to cell-free systems, *O. volvulus* larvae underwent two consecutive moults in the cell-based co-culture systems. The first moult (L3 to L4 = M1) was observed within 3–7 days of culture while the second moult (L4 to L5 = M2) was observed between day 48 and day 78, in which L4 larvae cuticle ecdysis to become L5 larvae (Fig 4). The use of cell lines as feeder layer triggered parasites second moult and it was consistently observed that amongst the *O. volvulus* larvae that achieved the first moult, if not all, a great proportion underwent a successful second moult. *O. volvulus* moulting rates ranged from 0 % (DMEM – 10 % NCS - LEC) to 99.2 % (DMEM – 10 % FBS - LLCMK2). The cell-based co-culture system that was the best to support parasite moulting from L3 to L5 was DMEM – 10 % FBS - LLCMK2 (M1 = 69.2±30 % and M2 = 69.2±30 %), though no statistically significant difference was observed as compared to DMEM – 10 % NCS - LLCMK2 (M1 = 57.8±30.7 % and M2 = 49.2±41%), DMEM – 10 % NCS - HC04 (M1 = 52.8±21.5 % and M2 = 52.8±21.5 %) and DMEM – 10 % NCS - HEK293 (M1 = 58.8±18.2 % and M2 = 57.0±15.3 %) except to DMEM – 10 % NCS – LEC (M1 = 1.7±4.7 % and M2 = 0.0±0.0 %) as shown (Fig 5).

**Fig 4.**
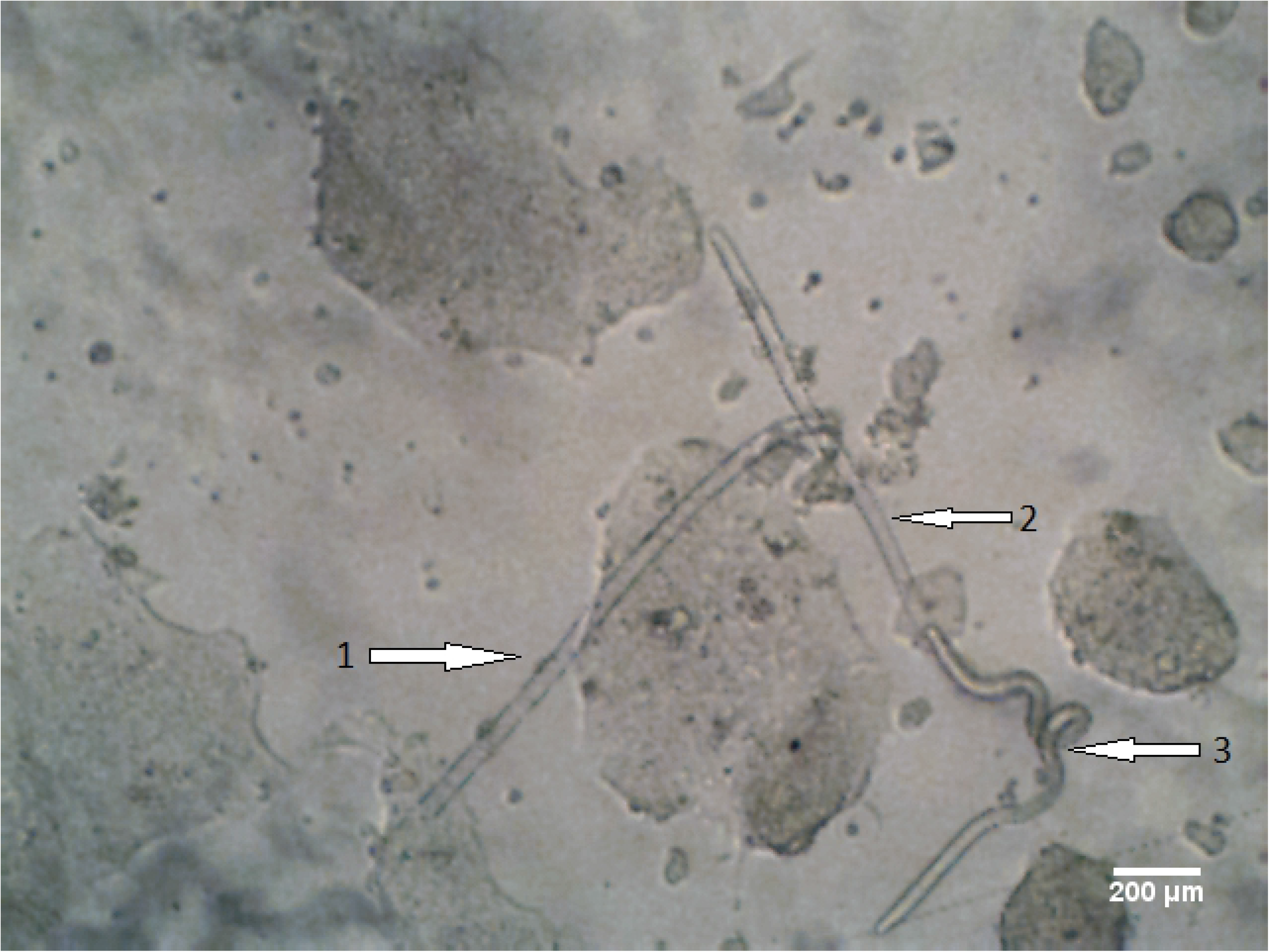
*O. volvulus* L4 larvae cuticle ecdysis to become L5 (1 = casted cuticles, 2 = cuticle being casted and 3 = newly moulted L5).

**Fig 5.**
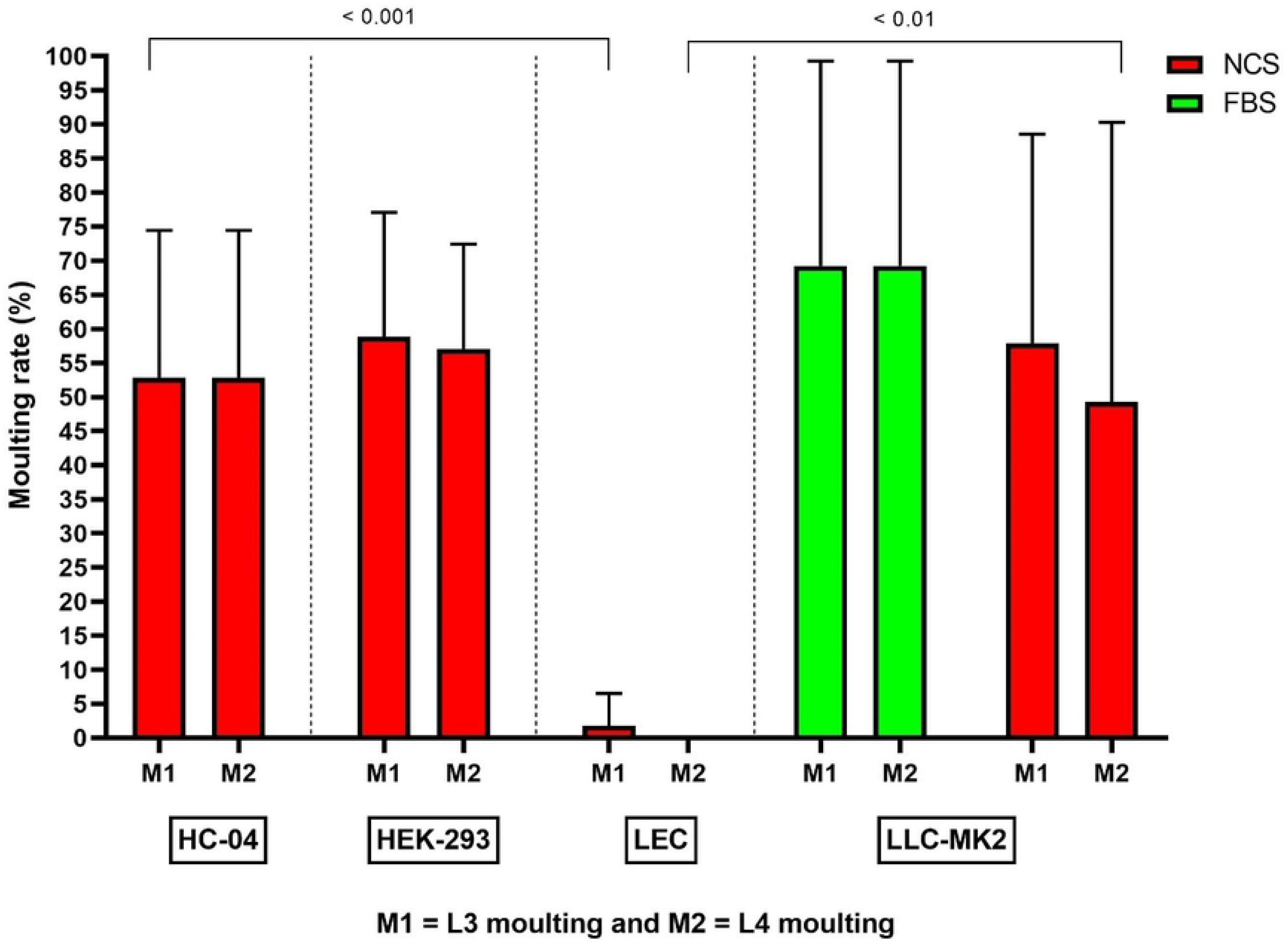
Influence of cell-based co-culture systems on *O. volvulus* larvae moults *in vitro.* (Results were pooled from different independent experiments, n =3 and each experimental setting conducted in quadruplets).

The length and width of *O. volvulus* larvae increased significantly throughout the duration of this study (233 days) and these changes were stage dependent. *O. volvulus* larvae morphometry increased as larvae switched from one stage to another. The L4 larvae length varied from 1450.17 μm to 2121.63 μm and 44.73 μm – 68.82 μm width, while L5 larvae length varied from 1478.09 μm to 3349.74 μm and 38.82 μm – 114.54 μm width. Although the length and width of both stages overlapped since male worms are shorter female worms, significant difference was observed between median length and width of L4 and L5 stages. Highest L5 larvae lengths were observed in NCS/LLC-MK2 (length = 3327.48 μm / width = 81.51 μm) and FBS/LLC-MK2 (length = 3349.74 μm / width = 114.54 μm) as shown in Fig 6 and Fig 7. Outlier points from either NCS/LLC-MK2 and FBS/LLC-MKS co-culture systems proved to be young female adult worms as worm mating was later observed.

**Fig 6.**
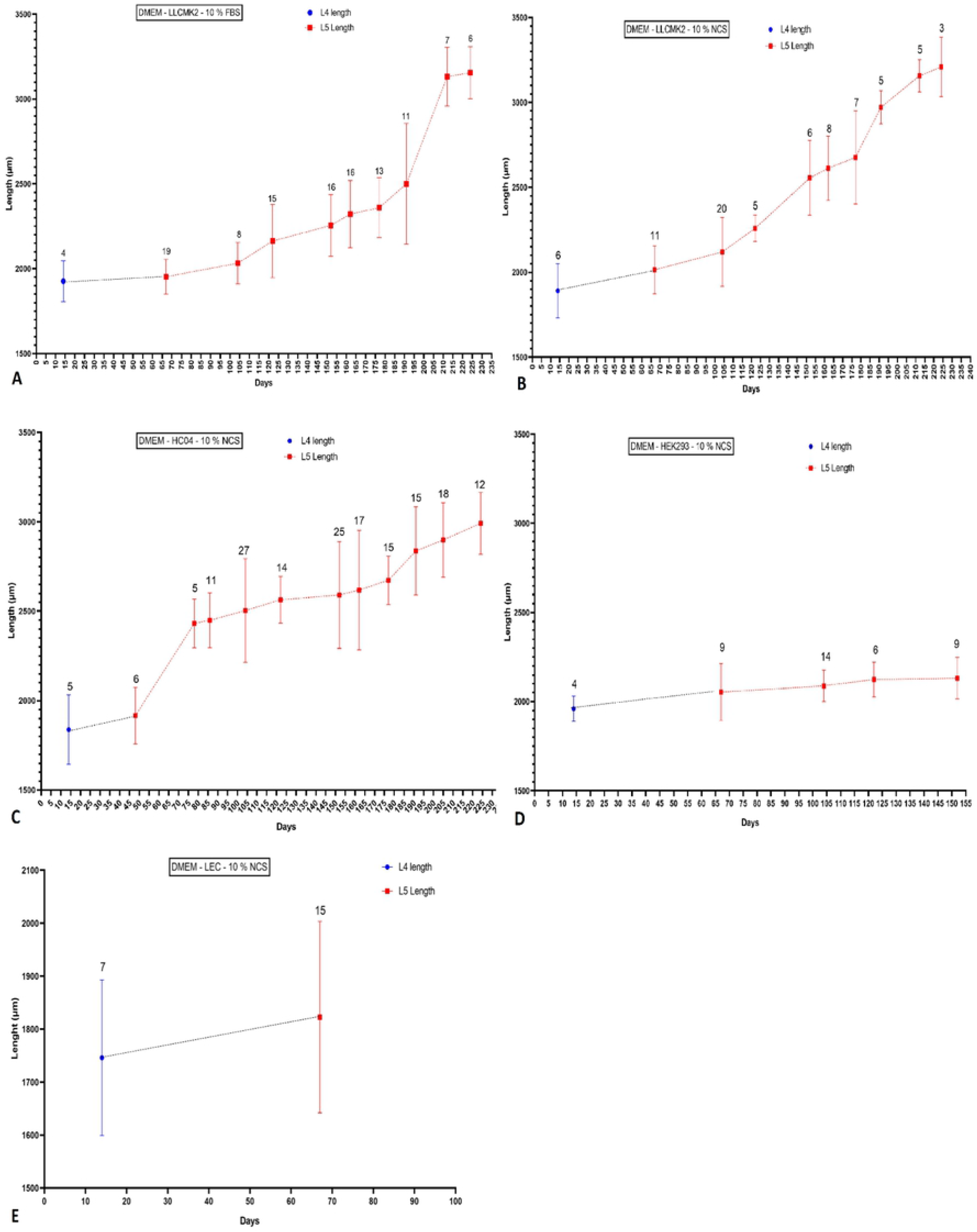
Length of *O. volvulus* L4 and L5 larvae observed from the *in vitro* cell-based co-culture systems with respect to days. (Bar representing the median). **A** DMEM – LLCMK_2_ – 10% FBS. **B** DMEM – LLCMK_2_ – 10% NCS. **C** DMEM – HC04 – 10% NCS. **D** DMEM – HEK293 – 10% NCS. **E** DMEM – LEC – 10% NCS.

**Fig 7.**
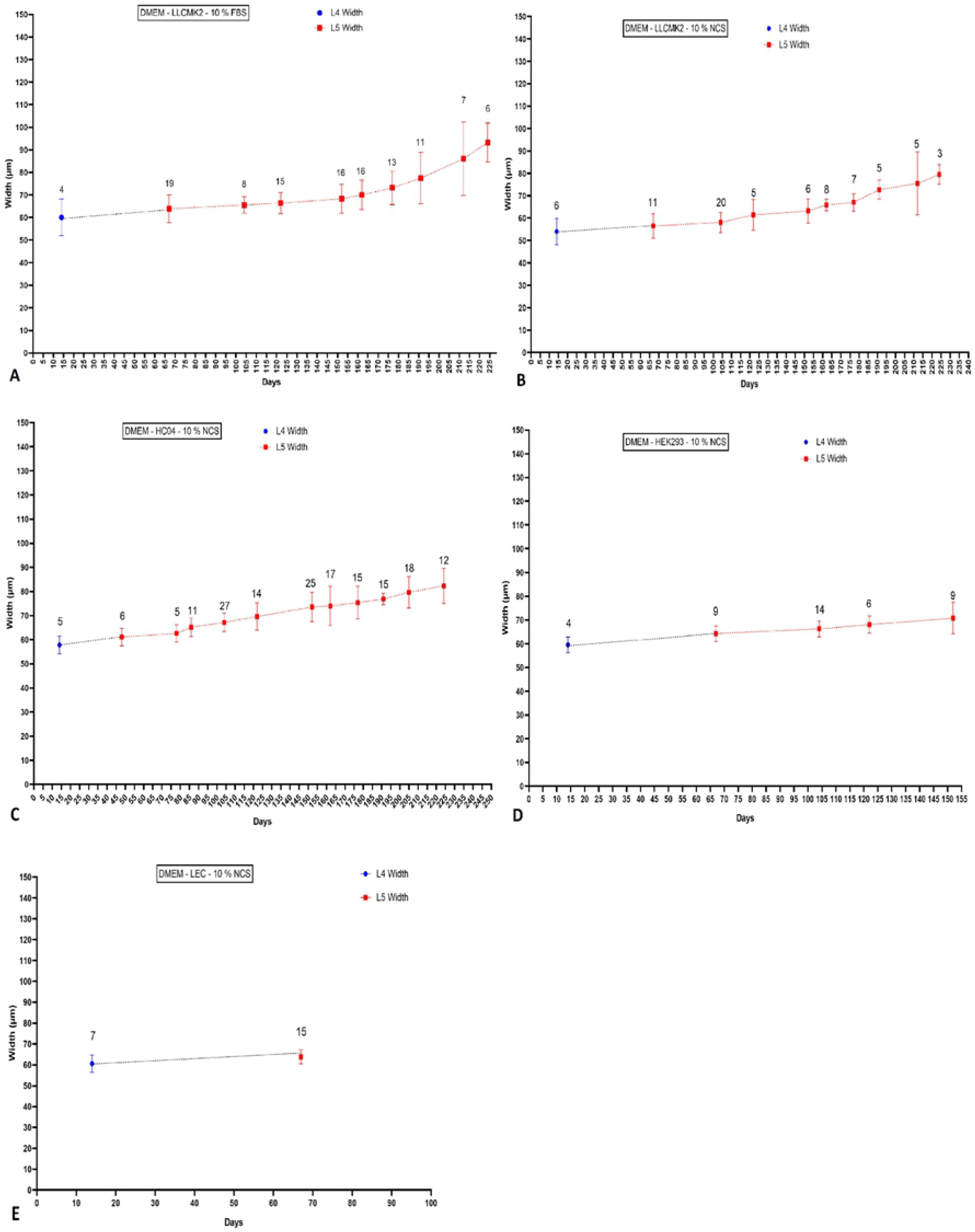
Width of *O. volvulus* L4 and L5 larvae observed from the in vitro cell-based co-culture systems with respect to days (Bar representing the median). **A** DMEM – LLCMK_2_ – 10% FBS. **B** DMEM – LLCMK_2_ – 10% NCS. **C** DMEM – HC04 – 10% NCS. **D** DMEM – HEK293 – 10% NCS. **E** DMEM – LEC – 10% NCS.

### Divergence in morphometry pattern of *O. volvulus* male and female worm’s

The length of female and male adult worms varied according to the cell-based co-culture system where they were cultured. Among all cell-based co-culture systems used, only DMEM – LLCMK_2_ – 10% NCS, DMEM – LLCMK_2_ – 10% FBS and DMEM – HC04 – 10% NCS displayed the best appreciable differences. Globally, no sex related *O. volvulus* larvae differentiation was possible between day 0 and day 104. Within this interval (0 – 104 days), the length of larvae recorded showed a wide variation of values but with no definitive conclusion on parasite sex. Above day 104, the discrepancy earlier observed in worm’s length values started dropping, therefore, leading to better categorization of worms based on their length. At most, *O. volvulus* adult male worms did not exceed 2900 μm while the female adult worms could reach up to 3300 μm (Fig 8). In addition to the differences in worm length, the early development of gonads of gonads was also used as discriminatory indicator between female and male adult worms.

**Fig 8.**
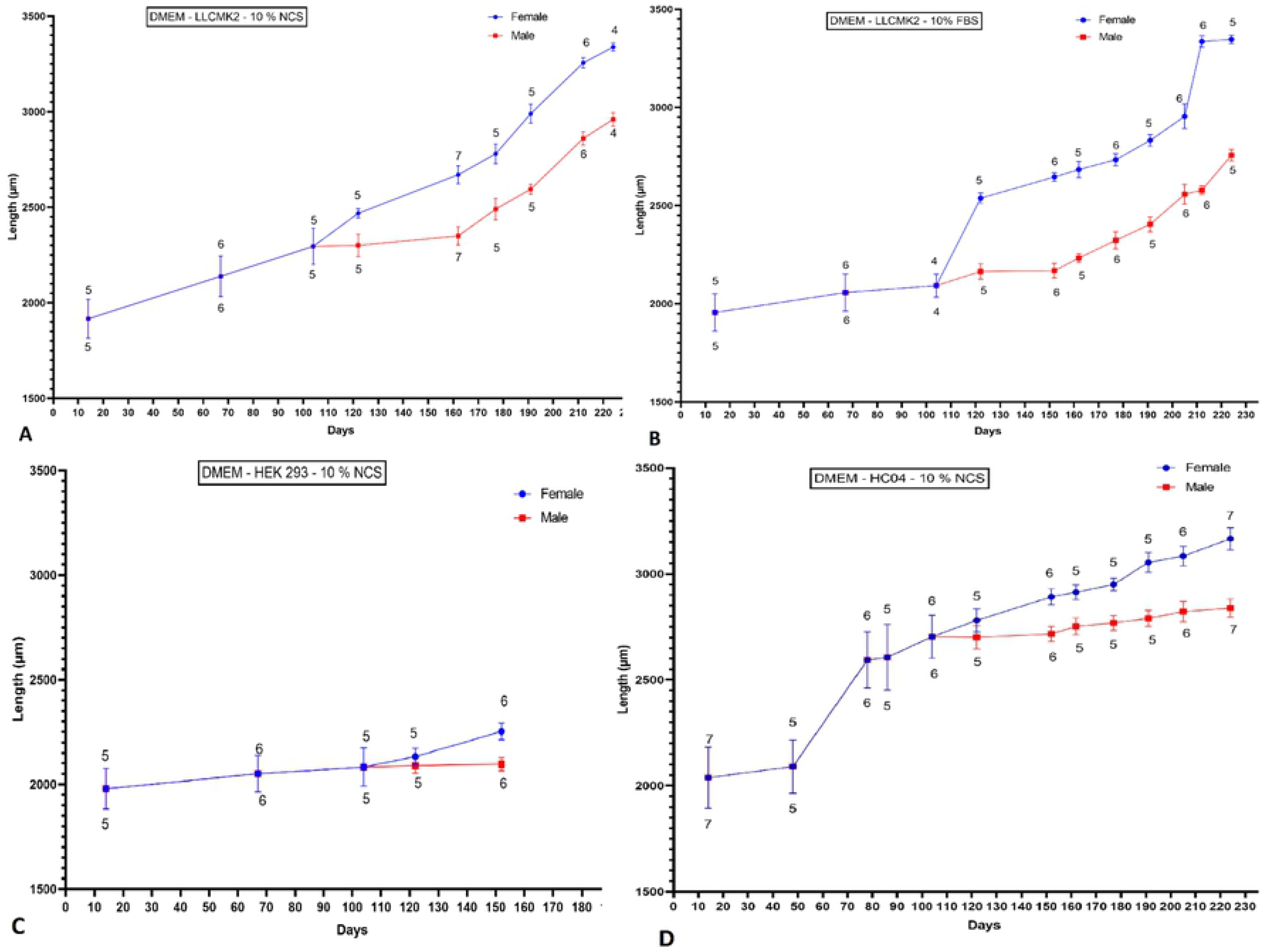
Disparity in sex-dependent worm morphometry changes in different cell-based co-culture systems. **A** DMEM – LLCMK2 – 10% NCS. **B** DMEM – LLCMK2 – 10% FBS. **C** DMEM – HEK293 – 10% NCS. **D** DMEM – HC04 – 10% NCS.

### Evidence of *O. volvulus* adult worms’ maturity: Mating and mating competition *in vitro*

Generally, experiments involving co-cultured systems were monitored for up to 233 days except for DMEM – 10 % FBS - LLCMK_2_ and DMEM – 10 % NCS - LLCMK_2._ The latter were the sole co-culture systems which could support the growth, development and mating of *O. volvulus* larvae from their infective larvae stage to adult worms and parasites were monitored in both systems for up to 315 days. Early L4 moults into L5 larvae were observed as from day 48 and lately by day 78. Newly moulted L5 larvae required an additional 4 months and 2 weeks (134 days) *in vitro* maintenance before the first parasite mating could be observed. Parasites copulation was observed only in the DMEM – 10 % FBS - LLCMK_2_ cell-based co-culture system. By day 212 of culture, the first *O. volvulus* adult worms mating was recorded (Figs 9A and B, S1 Media) and 12 days later a scene of mating competition was also recorded. The mating competition involved 2 adult males battling to copulate with an adult female worm (Fig 9C, S2 Media). The mating competition lasted for 11 days after which only one single adult male succeeded to mate with the female. The victorious adult male worm continued mating with the female worm for up to day 294. In summary, the whole mating process had a duration of 82 days and these copulated parasites survived for another 3 weeks (Figs 9D and F).

**Fig 9.**
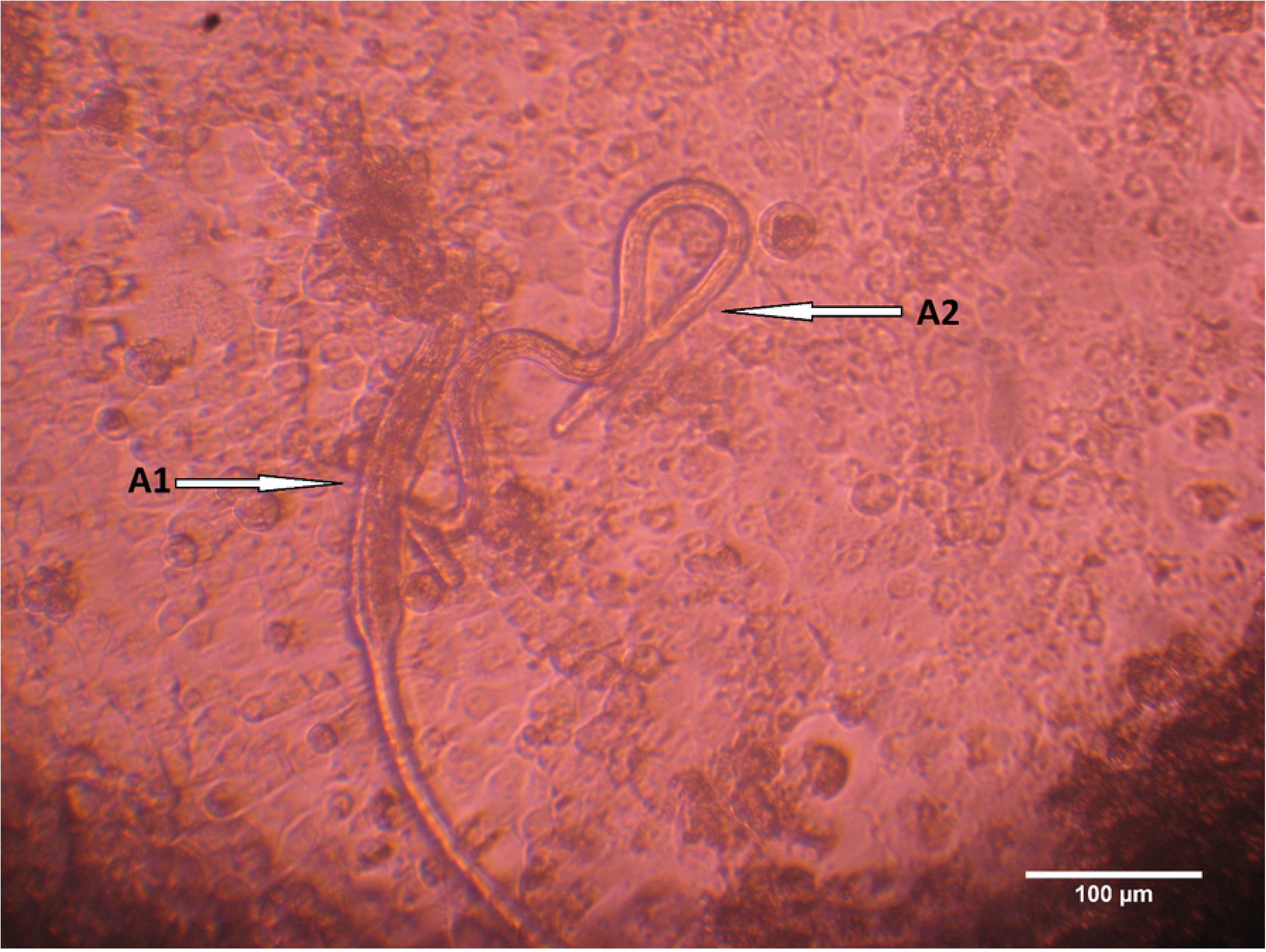
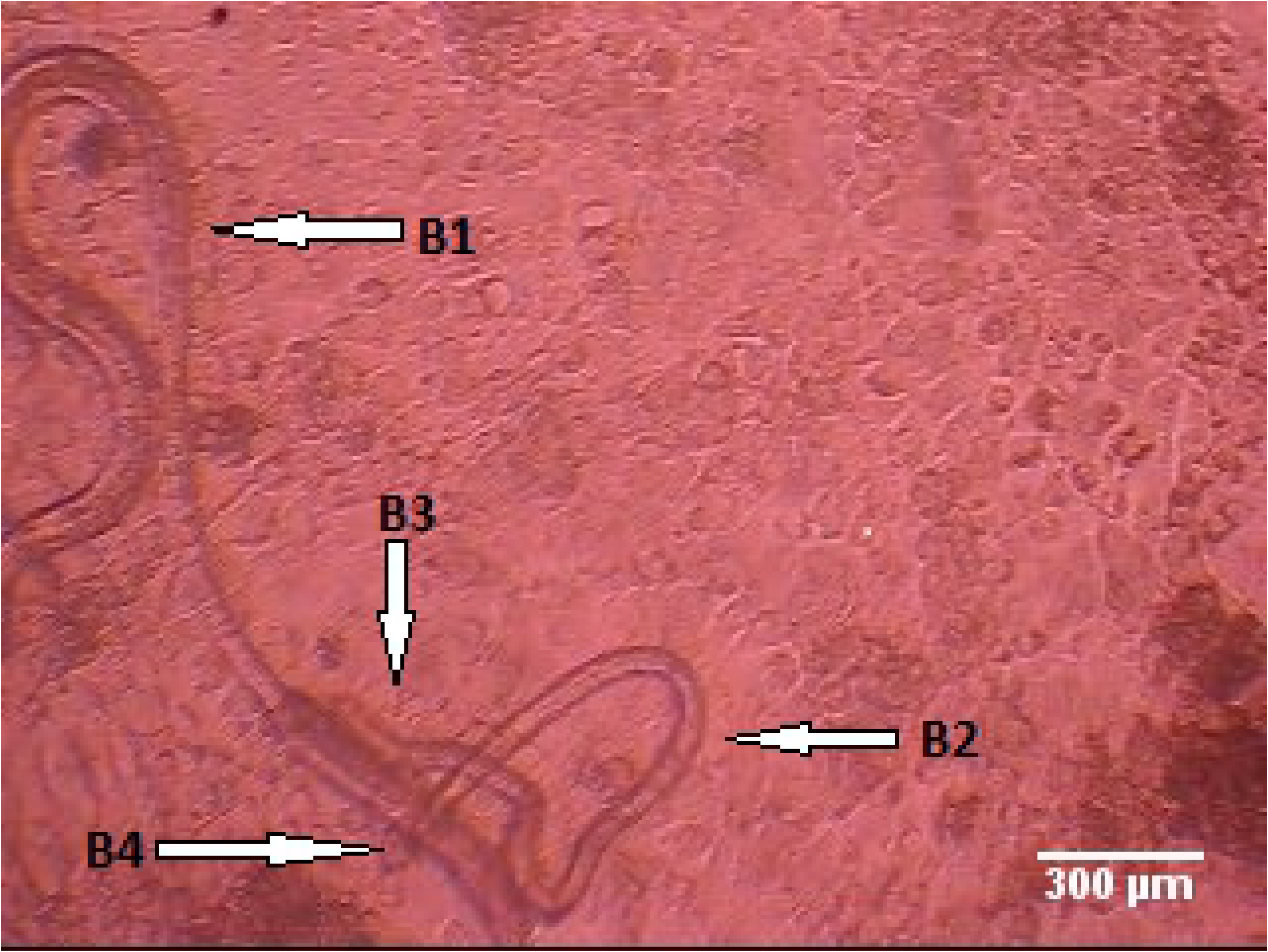
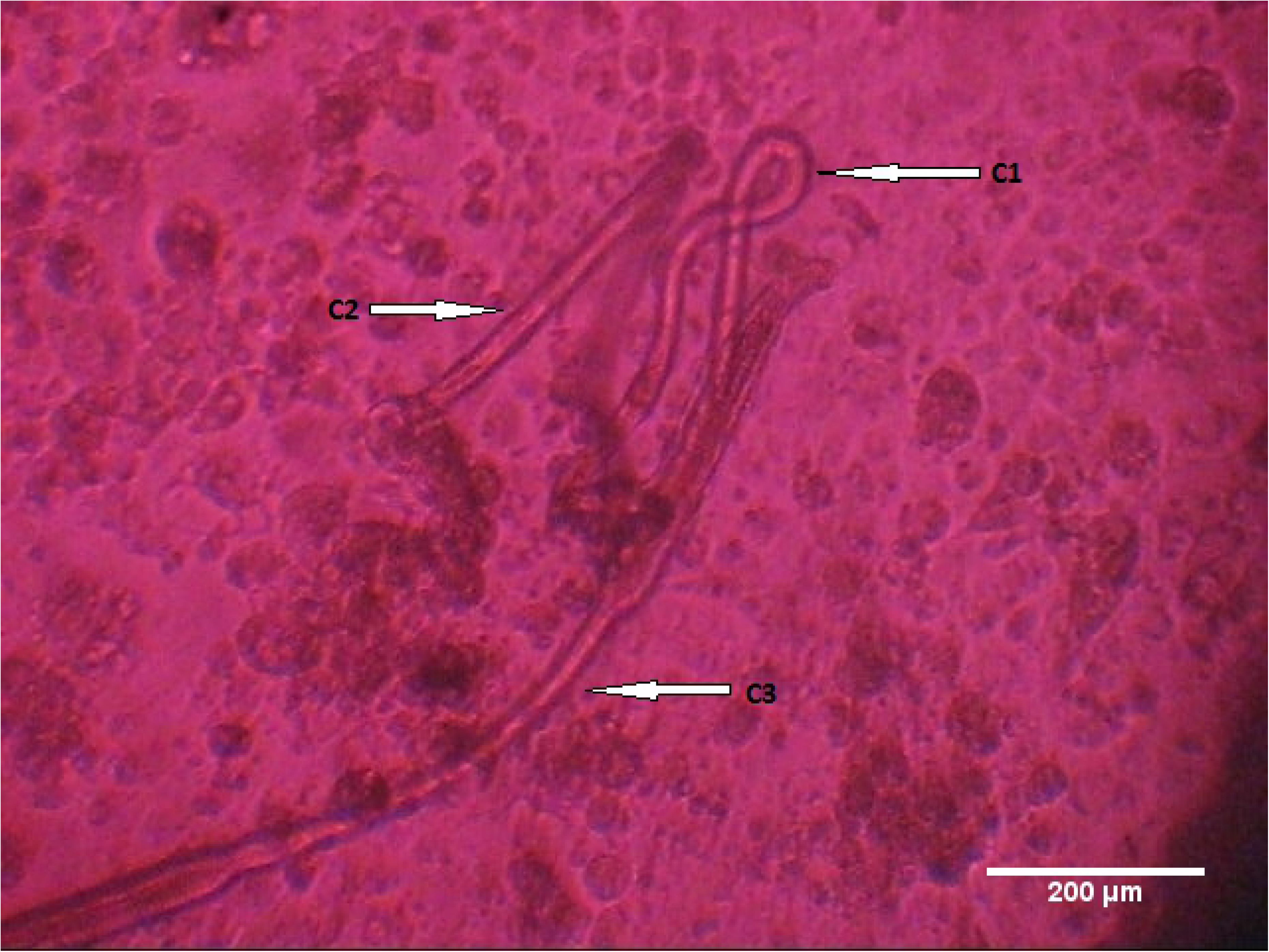
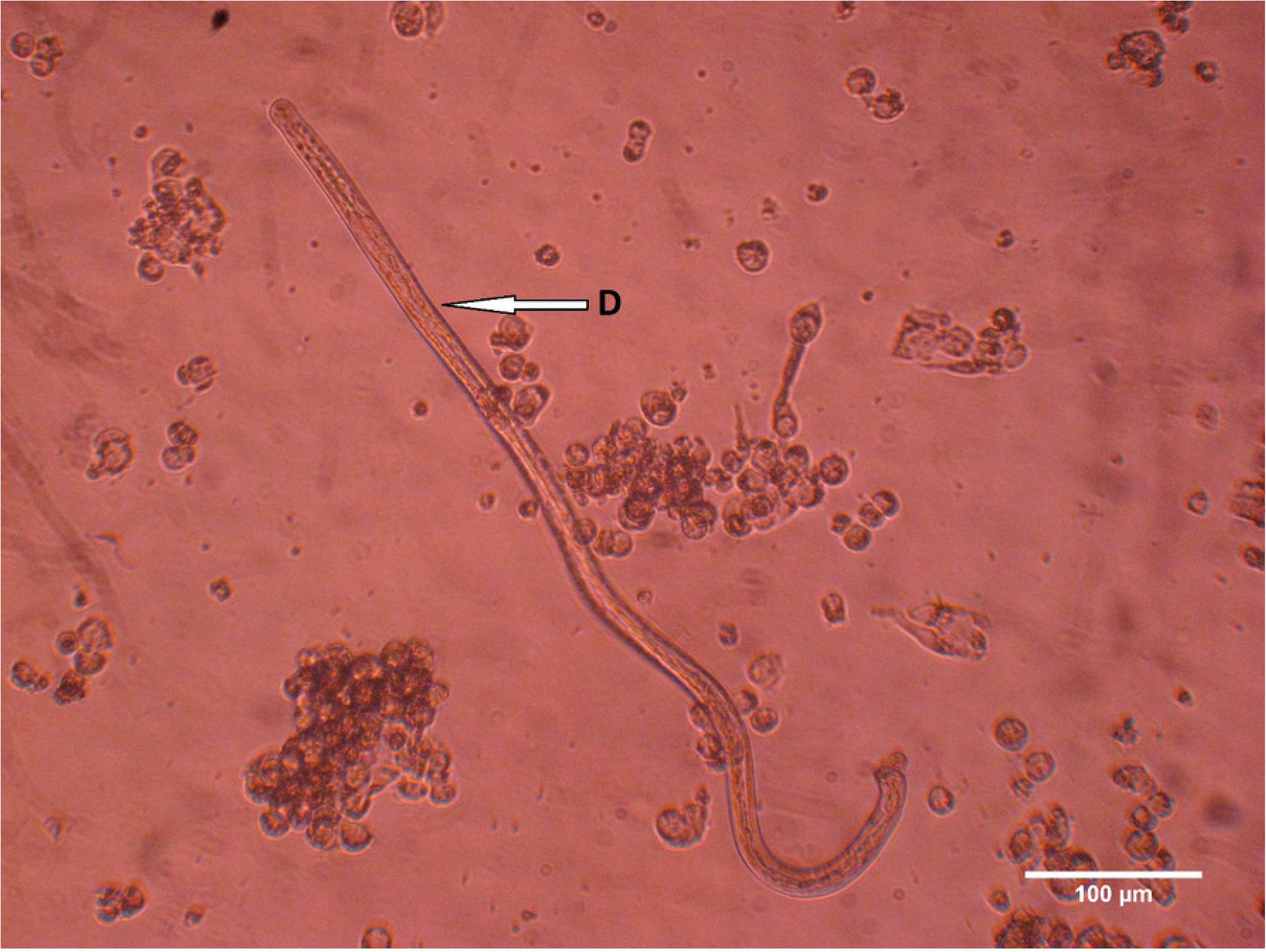
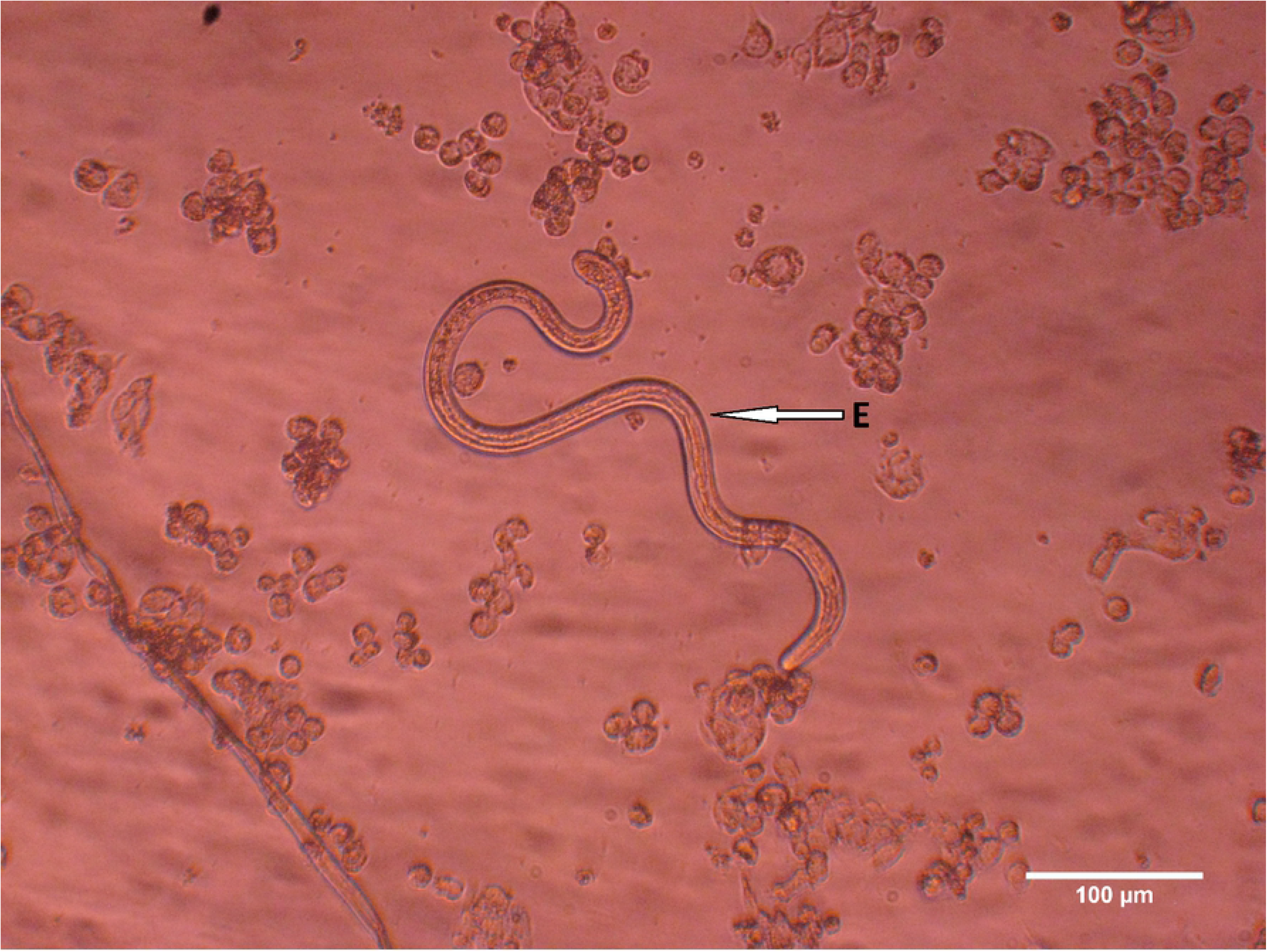
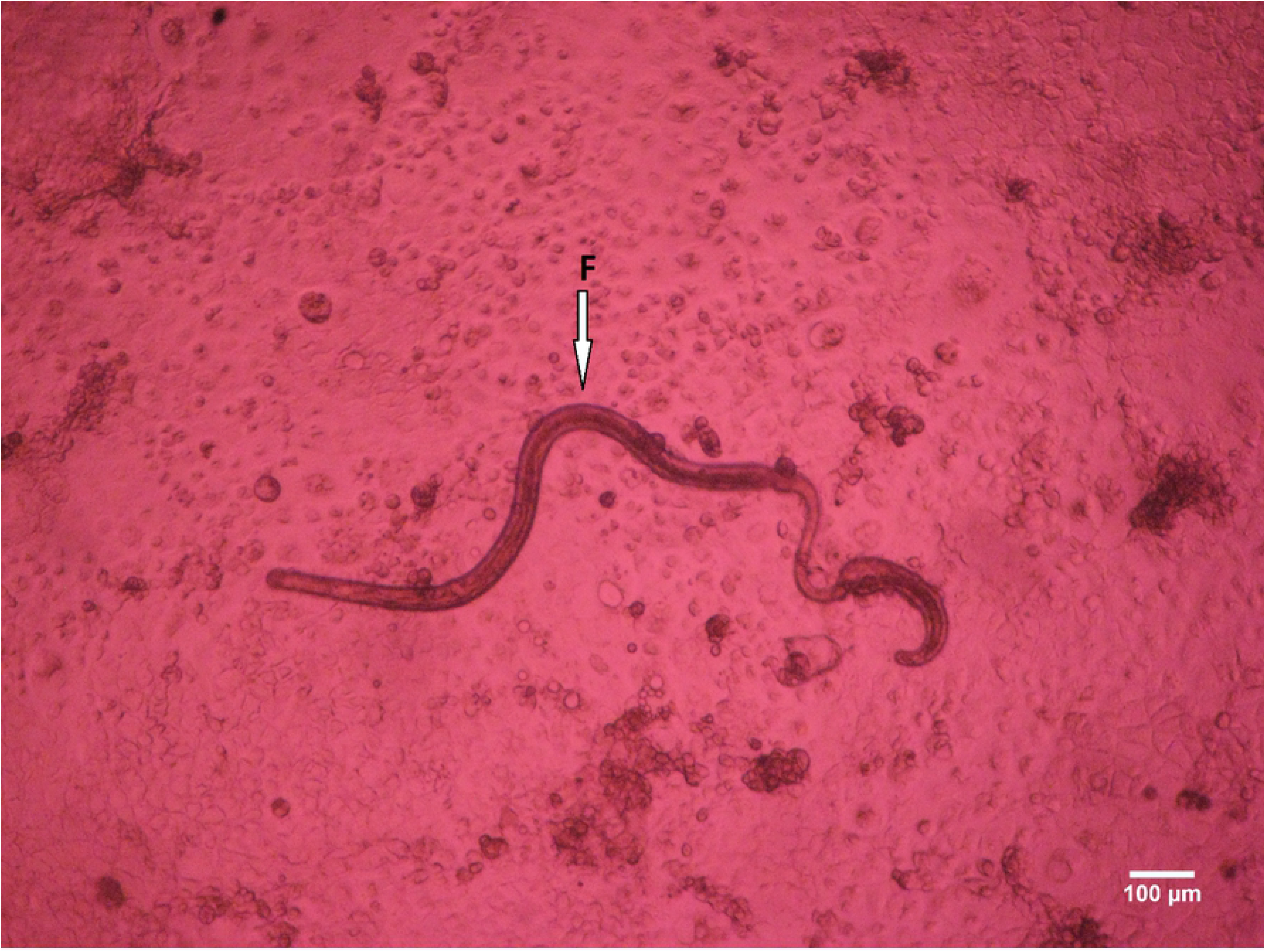
*O. volvulus* adult worms mating and mating competition. **A** Copulation topography during single mating. (A1: Posterior region of an adult female worm, A2: Anchored adult male worm mating with an adult femle). **B** Copulation topography during single mating. (B1: Mating adult female worm, B2: Mating adult male worm, B3: Copulation region entailing male specules insertion into female worm vulva, B4: Male and female worms anchor region).**C** Copulation topography during mating competition (C1: First adult male worm in competition for mating, C2: Second male worm in competion for mating, C3: Sollicited adult female worm for mating).**D** Adult female *O. volvulus* worm. **E** Adult male *O. volvulus* worm.**F** Mated adult female worm.

Overall, the co-culture system (DMEM – 10 % FBS - LLCMK_2_) could sustain and support *O. volvulus* larvae for up to 315 days and exhibiting parasite mating and mating competition. The proportion of adult worms involved in the mating and mating competition phenomenon is shown Table 1.

**Table 1.**
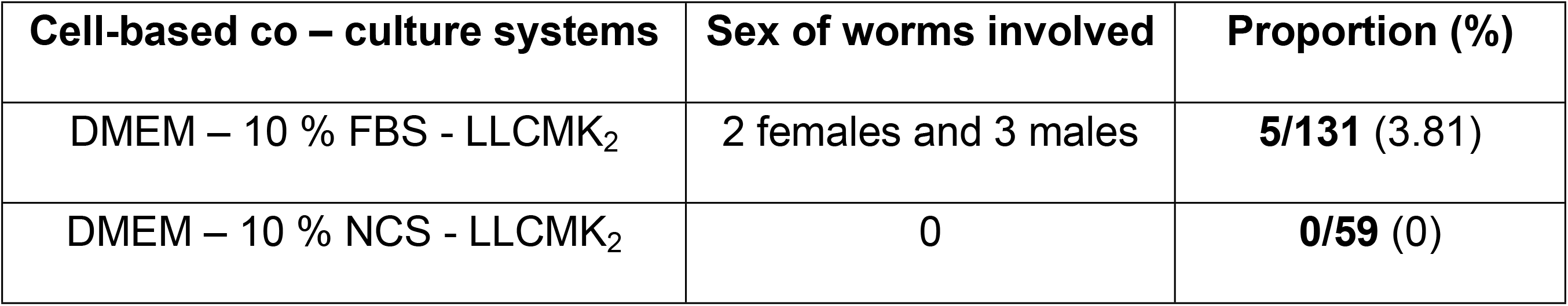
Proportion of adult worms involved in the mating process

### Cellular aggregation around *O. volvulus* adult worms in the *in vitro* cell-based co-culture system

From day 177, we observed changes in worm environment, notably cellular aggregation around the *O. volvulus* L5. This feature was displayed only in co-culture systems, mainly LLC-MK_2_ in combination either with DMEM – 10 % NCS or DMEM – 10 % FBS. The process started with the recruitment of thin transparent and insoluble cell-derived particles along worm cuticle (Figs 10A and B, S3 Media). These cell-derived transparent and insoluble particles lately came together to form a globular or oval shape aggregation around the worms that remained trapped in this mass (Figs 10C and D, S4 Media).

This process lasted for 138 days after which parasites perished. The proportion of adult worms involved in nodulogenesis is shown Table 2.

**Fig 10.**
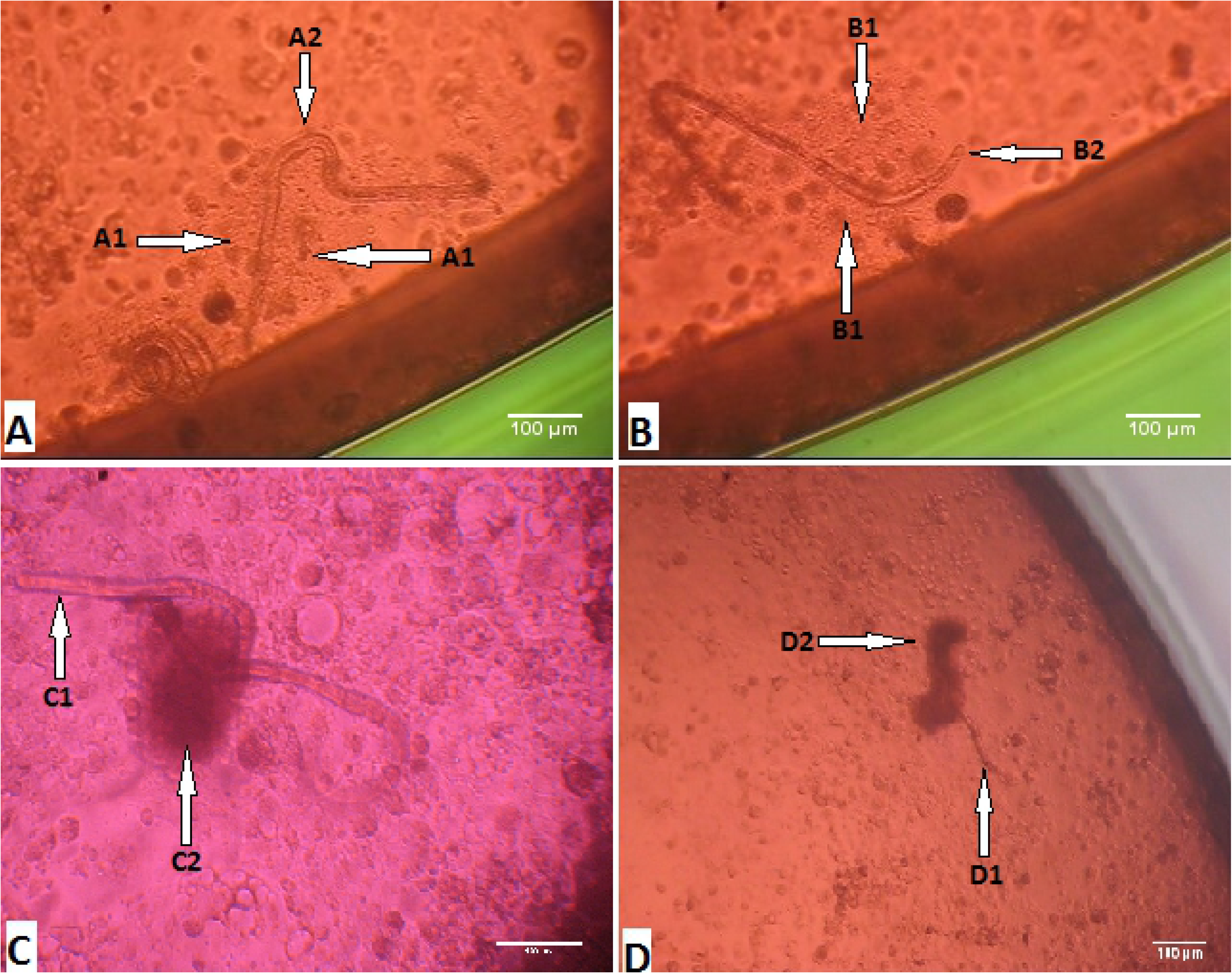
Cellular aggregation around *O. volvulus* young adults *in vitro*. **A&B**(A1 and B1: Early recruitment of cell-derived insoluble particles along the worm external wall, A2 and B2: Young adult worm trapped in the cellular aggregate). **C&D**(C2 and D2: Gathering of cell-derived insoluble particles into a globular/oval mass shape engulfing the young adult worms, C1 and D1: Young adult worm being engulfed).

### Linear regression analysis: factors influencing the growth and development of *Onchocerca volvulus larvae in vitro*

The contribution of the various culture media and supplements used in the improvement of worm viability were identified based on their standardized coefficient (S5 Table). Among these factors, the presence of feeder cells pre-eminently influenced *O. volvulus* larvae viability, HC-04 feeder cells were classified as topmost factor (β = 0.299) followed by LLC-MK_2_ feeder cells (β = 0.269), HEK-293 feeder cells (β = 0.129), LEC feeder cells (β = 0.060). Although the basic culture media types also promoted *O. volvulus* larvae viability, their impact was less important than that of feeder cells but higher as compared to that of sera supplements. DMEM basic medium had the leading effect (β = 0.042) and the least effect was observed with IMDM (β = − 0.068 10^-3^). Both sera supplements, FBS (β = − 0.074) and NCS (β = − 0.052) had unfavourable effect as compared to RPMI. The model was diagnosed by assessing the assumptions of normal distribution and homoscedasticity. The histogram of the residuals (errors) in the model was used to check if they are normally distributed (S6 Figure). Although not perfect, the frequency distribution of the residuals displayed a shape close to that of the normal Gauss curve, indicating evidence of normal distribution. Additionally, Q-Q plot was used for further check (S7 Figure). Here, the theoretical and observed quantiles were closed suggesting that the assumption of normal distribution of the residual was far to be not violated. The model was used to predict T_20_ and T_10_ values (Days) which correspond to the duration at which 20 and 10 % of the worms were still active (score 3) as shown in (S8 Figure). Co-culture with feeder cells in DMEM medium represented the systems that could extend survival of parasites for longer periods.

**Table 2.** Proportion of adult worms involved in nodulogenesis

| Co – culture systems | Number of adult worms involved | Proportion (%) |
| --- | --- | --- |
| DMEM – 10 % FBS - LLCMK<sub>2</sub> | 26 | **26/129** (20.15) |
| DMEM – 10 % NCS - LLCMK<sub>2</sub> | 10 | **10/59** (16.949) |

## Discussion

This study was conducted to establish an *in vitro* platform which could support and promote the growth and development of *O. volvulus* larvae from the *Similium* derived L3 larvae to the adult stages. Several attempts have been carried out in the perspective of achieving the complete life cycle of *O. volvulus* human stages in *in vitro* systems. Early studies used serum/cell free systems to culture filarial parasites and reported the maintenance of their full viability for up to a week [12–22]. Our previous reports and those from other investigators highlighted improvement of the culture conditions by supplementing the basic culture media used with serum or other synthetic additives. The serum-based culture systems achieved better parasite longevity and cuticle casting [21, 23–30, 32]. Due to challenges of inconsistency of serum composition, other researches opted to develop serum-free culture systems and co-culture systems using eukaryotic cells as feeder layers which yield best results obtained so far [33–43]. These studies did not provide a consensus on filarial parasite nutritional needs. Zofou *et al.* 2018 [42] established a hierarchical profile of most used *in vitro* culture supplements for filarial parasites, in which the presence of feeder cells was ranked as most top important followed by culture supplementation with serum and finally the culture media type. Based on these observations, four feeder layer cell line types (LLC-MK_2_, HC-04, HEK-293 and LEC), two serum types (FBS and NCS) and five basic culture media (RPMI-1640, DMEM, MEM, NCTC-135 and IMDM) were evaluated either in combination in the cell-free systems or in the co-culture systems for the growth and development of *O. volvulus* larvae *in vitro*.

In order to assess *O. volvulus* larvae *in vitro* growth and development five variables were monitored: mean motility, moulting rate, parasite stage specific morphometry, mating and nodule formation. With respect to *O. volvulus* larvae viability, parasite mean motility was used as first indicators among others (mean motility, moulting rate, worm morphometry). Regardless the culture system, *O. volvulus* larvae motility displayed a ‘‘saw teeth’’ evolution pattern. For both systems (cell-free and co-culture systems), cultured L3 larvae motility started reducing as from day 0 and persisted still day 3 when the first L3 moults were observed. The switch from one stage to another was in first instance marked by a significant drop in motility followed by an abrupt increase in motility (shift from score 2 to score 3), and this same phenomenon was also later on observed when larvae moulted from L4 to L5. The progressive drop in motility till moulting followed by a drastic increase could be an evidence that the greatest fraction of energy produced by the larvae is stilted towards cuticle ecdysis which is prioritized at this time point and less energy assigned to worm twitching. Page and Johnstone [48] reported that the moulting process in *C. elegans* nematodes is preceded by a period of decreased general activity and feeding, known as lethargus, when the old cuticle begins to disconnect from the underlying hypodermis. Singh and Sulston [49], stated that during apolysis, the old cuticle is separated, allowing the newly synthesized one to be secreted in the space between the two layers. The moulting cycle is completed with ecdysis, when the old cuticle is completely shed and the worm emerges to the next stage with a new cuticle. Moreover, it was observed that, only co-culture systems could support *O. volvulus* larvae moulting from L4 to L5. These are further indications ascertaining that some nutrients secreted/excreted by feeder cells were key factors for parasite development as it was clearly demonstrated that no L4 to L5 larvae moults were observed in cell-free culture systems. Toback *et al.* [50] reported that kidney epithelial cells release growth factors such as epidermal growth factor (EGF), transforming growth factor-type - alpha (TGF-α), insulin-like growth factor I (IGF-I), platelet-derived growth factor (PDGF), and insulin which are being exploited by the parasite for their growth and development. McConnell *et al.* [51] also showed that non-transfected HEK-293 release nerve growth factor which were beneficial to *O. volvulus* larvae growth and development. Without feeder cells, *O. volvulus* larvae stayed viable *in vitro* for up to 84 days, their longevity rose by 3.7 folds in the co-culture system (315 days). The co-culture system developed in this study is superior to the one reported by Voronin *et al.* [44] in the sense that they used two distinct culture settings to achieve L5 stage while our system made use of a single cell – based co-culture system (DMEM – LLC-MK2 – 10% NCS/FBS) and the maximum attainable longevity they reported for their system was 117 days versus 315 days for this new system. Additionally, our system achieved higher L3 and L4 moulting rate (69.2±30 %) as compared to theirs (Max. of 60 %). Moreover, our single cell-based co-culture system supported adult *O. volvulus* worms mating, mating competition and early nodulogenesis that wasn’t achieved with their system. Our findings open up new avenues for drug screening and in-depth investigation of *O. volvulus* biology.

Moulting entails synthesis of the new skin and shedding of the old, and represents an important phenomenon for the growth and maturation of filarial parasites. The moulting process is critical for filarial parasites and disruption of moult can have serious consequences for survival and reproductive success. It is vital for filarial parasites to undergo two consecutive moults and metamorphosis in order to become fully mature. The cell-free culture system could only support the first moult (M1) of *O. volvulus* infective larvae leading to L4 stage larvae ranging from 0 % (MEM – 10 % NCS) to 78.8±13.2 % (DMEM – 10 % NCS), the basic culture medium type combined with animal serum used provided the necessary nutrients (proteins, electrolytes and hormones) required by *O. volvulus* infective larvae to moult to L4. The second parasite moult (M2) was only observed in the co-culture system. It was therefore clearly established that the feeder cells play a crucial role in the development and maturation of *O. volvulus* parasite. Previous studies to culture other filarial parasites also demonstrated the pre-eminent role of feeder cells in their successful *in vitro* maintenance [26, 41, 42, 44]. Except from (DMEM – 10 % NCS – HEK293), (DMEM – 10 % BCS – LLC-MK2) and (DMEM – 10 % NCS – LLC-MK2) co-culture systems, all *O. volvulus* L3 larvae that successfully undertook the M1 moults equally performed the M2 moults. Highest M1 and M2 moulting rate were reported in (DMEM – 10 % FBS – LLC-MK2), which were 69.2±30.0 % and 69.2±30.0 %, respectively. This could account to the fact that FBS has served as an additional source of essential nutrients, growth factors and hormones as the monkey kidney cells. Moreover, FBS could bind and protect essential nutrients that are otherwise unstable. It could also function to neutralize toxic substances in the medium or supply necessary transport factors or enzymes [52].

*O. volvulus* morphometry was also used as an indicator to assess parasite growth, although *O. volvulus* L4 stage morphometry overlapped with the L5 stages. The length of L4 stages ranged from 1450 μm to 2122 μm (Length median = 1794 μm) and width from 45 μm to 69 μm (Width median = 57 μm) and L5 length ranged from 1478 μm to 3350 μm (Length median = 1892 μm) and width from 39 μm – 115 μm (Width median = 60 μm). There was no significant difference between L4 larvae measured from cell-free systems and L4 larvae from co-culture systems that failed to moult to L5 stages, whereas L4 larvae lengths significantly differ from those of L5. Highest L5 larvae length was recorded in (DMEM - 10 % NCS – NCS) and (DMEM – 10 % FBS – LLC-MK2) co-culture systems, moreover L5 larvae length from both systems differed significantly (*P =* 0.0418). We also noticed that outlier lengths recorded in (DMEM - 10 % NCS – NCS), (DMEM – 10 % FBS – LLC-MK2) and (DMEM - 10 % NCS – HC-04) co-culture systems were female *O. volvulus* larvae, which were later involved in mating and mating competition.

During this study, evidences of adult worm maturity were given. In filarial parasite biology, mating can only occur when parasites are mature with well-developed gonads. Moreover, these parasites have to be mature enough to be able to produce and respond to mating related signals for the process to be carried on. For the first time ever, *O. volvulus* adult parasites mating behaviour was documented and photographed. The first mating scene was observed as from day 212 as *O. volvulus* larvae were co-cultured in (DMEM – 10 % FBS – LLC-MK2). Within 12 days, two adult male worms battled/competed to copulate with a female. At the end of the process, only the victorious male worm succeeded to copulate with the female worm. As earlier reported with *C. elegans in vitro* [53–55], the initiation of *O. volvulus* female worm copulation with a mature male worm may had been triggered by attractants (hormones) excreted/secreted by the ready-to-mate female worm. The presence of this chemotactic substance may have at first instance attracted the first adult male worm then the second which resulted to a fierce struggle for mating. The nature and composition of this attractant is unknown and requires extensive investigations. In total, the mating process took 82 days after which these parasites remain viable for 3 more weeks (21 days). Trees *et al.* 2000 reported that it takes 279–532 days post infection (dpi) for the closely related *O. ochengi* parasite of cattle to develop into fully mature and fertile adult worms capable of releasing microfilariae and more than 400 days post infection (dpi) for *O. volvulus* to do the same in a chimpanzee model [8, 11]. Since the female worms survived only 3 weeks after copulation, we did not observe release of microfilariae in this system. It’s possible that at this time point, the female worms needed a particular stimulus either from the environment or self-produced to trigger the embryogenesis and later on the release of microfilariae. This calls for further investigation to generate the complete reproduction cycle of *O. volvulus in vitro.*

*Onchocerca volvulus* adult worms that were involved in both the mating process followed by formation of nodules survived for a longer period of time (315 days) as compared to those that failed to undergo these events (234 days). In the chronology of biological events, *O. volvulus* adult worms recruited of cell derived insoluble particles as early as from day 177 which was later exhibited by a linear aggregation of these particles along the parasite. The next event entailed parasites mating which occurred as from day 212 and finally these cell-derived insoluble particles gathered into a globular/oval shape mass engulfing the concerned parasites as from day 224. This event could indicate early nodulogenesis, however further studies are required to clarify the composition of these aggregates as well as its role in worm development (an attempt for the parasite to produce a shelter to protect itself so that it can complete its developmental cycle as observed with nodule formation in the mammalian host). According to Collins *et al.* [56] nodule forms only around female worms and mating probably occurs before or early during nodule formation. The production of microfilariae by the female *O. volvulus* is not essential for nodule formation since many nodules contained non-fecund, living females. Al-Qaoud et al. [57] reported that filarial worm encapsulation in the murine model was IL-5 dependent. These observations on the incomplete nodule formation *in vitro* deserves further investigations by providing to the *in vitro* system some immunologic effectors that exist *in vivo.*

## Conclusions

This study has successfully established an *in vitro* platform for *O. volvulus* growth and development that mimic the parasite biology in the human host. The platform enables us to culture *O. volvulus* for 315 days and observe for the first-time moulting (L3-adult worms/L5) and mating behaviour as well as mating competition and early phase of nodule formation. The establishment of this platform therefore stands as an important achievement in *O. volvulus* developmental biology and has potential for the identification of targets for drug discovery against different phases of development of this filaria parasite.

## Abbreviations

APOC: African Programme for Onchocerciasis Control
OCP: Onchocerciasis Control Programme
PTS: Post-Treatment Surveillance
HC-04: Human Hepatocyte cells
HEK-293: embryonic human kidneys cells
LLC-MK2: Monkey Kidney cells
LEC: Mouse embryonic lung cells
L3: Infective Larvae
L4: Stage 4 larvae
L5: Stage 5 larvae
RPMI: Roswell Park Memorial Institute
DMEM: Dulbecco’s Minimum Essential Medium
MEM: Minimum Essential Medium
IMDM: Iscove’s Modified Dulbecco Medium
NCTC: New jersey Cell Type Collection
NCS: New-born Calf Serum
FBS: Foetal Bovine Serum
BCS: Bovine Calf Serum
MDA: Mass Drug Administration
WHO: World Health Organisation
CMFL: Community Microfilariae Loads
EGF: Epidermal Growth Factor
TGF-α: Transforming Growth Factor - alpha
IGF - I: Insulin-like Growth Factor – I
PDGF: Palettes-Derived Growth Factor
DDT: Dichlorodiphenyltrichloroethane.

## Acknowledgements

We thank Mr. Oben Bruno for his assistance in obtaining the necessary vector material for this work.

## Availability of data and materials

The datasets used and/or analysed during the current study are available from the corresponding author upon reasonable request.

## Competing interests

The authors declare that they have no competing interests.

## Funding

This work was funded through grants awarded to AH, MR, MPH and SW from the German Research Foundation (DFG) (Grant DFG HO 2009/10-1, HO 2009/14-1, HU 2144/3-1) within the “German-African Cooperation Projects in Infectiology”. In addition, SW and AH are financially supported by the Federal Ministry of Education and Research (BMBF, initiative Research Networks for Health Innovations in sub-Saharan Africa: TAKeOFF) as well as SW, MPH and AH by the Horizon 2020 Framework Programme: HELP. AH is a member of the Excellence Cluster Immunosensation (DFG, EXC 1023) and the German Center of Infectious Disease (DZIF).

## Author’s contributions

Conceptualization: AH, PAE, MPH, SW

Data curation: AJN, NVTG, SW

Formal analysis: AJN, NVTG, SW

Funding acquisition: AH, MR, MPH, SW

Investigation: NVTG, AJN, EJG, FFF, CAK, WPCN

Methodology: AJN, NVTG, MR, MPH, AH, SW

Resource provision: AH, MR, MPH, AH

Writing original draft: NVTG, AJN

Writing review and editing: AJN, NVTG, MR, MPH, MEE, SW

## Supporting information

**S1 Media.***O. volvulus* adult parasites mating *in vitro.*

**S2 Media.***O. volvulus* adult parasites mating competition *in vitro.*

**S3 Media.** Early recruitment of cell-derived insoluble particles *in vitro.*

**S4 Media.***O. volvulus* adult parasites early nodulogenesis *in vitro.*

**S5 Table.** Experimental factors introduced in the model influencing *O. volvulus* larvae viability and their standardized coefficients.

**S6 Figure.** Histogram of the residuals (errors) in the model for normal distribution and homoscedasticity.

**S7 Figure.** Q-Q plot suggesting normal distribution of the residual.

**S8 Figure.** Model predicted T_20 and_ T_10_ values (Days).

